# Increasing neural network robustness improves match to macaque V1 eigenspectrum, spatial frequency preference and predictivity

**DOI:** 10.1101/2021.06.29.450334

**Authors:** Nathan C. L. Kong, Eshed Margalit, Justin L. Gardner, Anthony M. Norcia

## Abstract

Task-optimized convolutional neural networks (CNNs) show striking similarities to the ventral visual stream. However, human-imperceptible image perturbations can cause a CNN to make incorrect predictions. Here we provide insight into this brittleness by investigating the representations of models that are either robust or not robust to image perturbations. Theory suggests that the robustness of a system to these perturbations could be related to the power law exponent of the eigenspectrum of its set of neural responses, where power law exponents closer to and larger than one would indicate a system that is less susceptible to input perturbations. We show that neural responses in mouse and macaque primary visual cortex (V1) obey the predictions of this theory, where their eigenspectra have power law exponents of at least one. We also find that the eigenspectra of model representations decay slowly relative to those observed in neurophysiology and that robust models have eigenspectra that decay slightly faster and have higher power law exponents than those of non-robust models. The slow decay of the eigenspectra suggests that substantial variance in the model responses is related to the encoding of fine stimulus features. We therefore investigated the spatial frequency tuning of artificial neurons and found that a large proportion of them preferred high spatial frequencies and that robust models had preferred spatial frequency distributions more aligned with the measured spatial frequency distribution of macaque V1 cells. Furthermore, robust models were quantitatively better models of V1 than non-robust models. Our results are consistent with other findings that there is a misalignment between human and machine perception. They also suggest that it may be useful to penalize slow-decaying eigenspectra or to bias models to extract features of lower spatial frequencies during task-optimization in order to improve robustness and V1 neural response predictivity.

**Author summary:** Convolutional neural networks (CNNs) are the most quantitatively accurate models of multiple visual areas. In contrast to humans, however, their image classification behaviour can be modified drastically by human-imperceptible image perturbations. To provide insight as to why CNNs are so brittle, we investigated the image features extracted by models that are robust and not robust to these image perturbations. We found that CNNs had a preference for high spatial frequency image features, unlike primary visual cortex (V1) cells. Models that were more robust to image perturbations had a preference for image features more aligned with those extracted by V1 and also improved predictions of neural responses in V1. This suggests that the dependence on high-frequency image features for image classification may be related to the image perturbations affecting models but not humans. Our work is consistent with other findings that CNNs may be relying on image features not aligned with those used by humans for image classification and suggests possible optimization targets to improve the robustness of and the V1 correspondence of CNNs.

## Introduction

Our visual system has the seemingly effortless ability to extract relevant features from the environment to support behaviour. Computational models known as convolutional neural networks (CNNs) incorporate principles of the neurobiology of the visual system and have allowed us to mimic some capabilities of our visual system [1]. These models have had immense success in artificial intelligence and can be trained to perform at or above human capabilities on many tasks in the visual domain such as object categorization and semantic segmentation [2–6]. This led to many comparisons between the internal representations of CNNs to those of the human and non-human primate ventral visual stream, showing that task-optimized CNNs are also quantitatively accurate models of visual processing [7–14].

Although these models show remarkable similarities to the primate ventral visual stream, they diverge significantly from humans in their classification behaviour on images that have been modified by human-imperceptible, non-random image perturbations. In particular, these *adversarial perturbations* can cause the model to completely mis-classify the image even though it could correctly classify the unperturbed image, resulting in poor *adversarial robustness* [15]. This is clearly an issue in safety-critical applications (e.g., self-driving cars), so the machine learning community has been developing techniques to train these models to be more robust to adversarial perturbations [16–21]. These robust optimization techniques have been shown to be able to defend against very strong adversarial attacks, although there still exist perturbations that can fool models trained with these techniques.

This striking misalignment between machine and human image classification on adversarially perturbed images suggests that humans are using image features for the task that are different from those used by models explicitly optimized to perform the task [22–25]. This leads to the following question: what are some properties of the internal representations that differ between primate and machine vision and that result in such brittleness?

Motivated to understand the dimensionality of the population code, Stringer et al. [26] developed a theory connecting the eigenspectrum of a system’s neural responses to the system’s vulnerability to small stimulus perturbations. In mouse primary visual cortex (V1), Stringer et al. [26] showed that the eigenspectrum of the neural responses to natural scenes decays according to a power law with exponent α ≈ 1. Their theory predicted this power-law-like behaviour and states that if *α* < 1 for neural responses to natural scenes, then the neural code is “pathological” in the sense that small perturbations in the stimulus could result in unbounded changes in the neural responses. Too much variation in the responses with respect to the stimulus would allow minute stimulus changes to drastically affect the neural responses. As the existence of adversarial examples in CNNs is, by definition, vulnerability of CNNs to small input perturbations, we investigated the eigenspectra of the representations of models that are either robust or not robust to these perturbations.

The eigenspectrum can also provide insight into the image features extracted by a system. Lower principal components are associated with neural response variance related to coarser stimulus features and higher principal components are associated with variance related to finer stimulus features (see Extended Data Fig 6 in Stringer et al. [26]). Thus, if one system’s eigenspectrum decays slower (i.e., has a smaller power law exponent) than that of another system, it means that a larger amount of neural response variance is dedicated to the encoding of fine stimulus features in the first system than that of the second system. We therefore hypothesized that model responses with eigenspectra of small power law exponents have many artificial neurons tuned to image features of high spatial frequencies.

To test the hypothesis that many artificial neurons are tuned to high spatial frequencies, we investigated the preferred spatial frequency tuning distributions of these models and compared these distributions to that of cells in the foveal area of macaque V1. This resulted in three main contributions. Firstly, we found that models with higher adversarial robustness have internal representations whose eigenspectra decay slightly faster than those of their non-robust counterparts and is consistent with the theory of Stringer et al. [26]. Secondly, by performing in-silico electrophysiology experiments, we found that non-robust models had a large proportion of neurons tuned to high spatial frequencies. Moreover, the similarity between a model’s preferred spatial frequency distribution and that of cells in the foveal area of macaque V1 was higher for robust models than that of non-robust models. Robust models, however, still had many artificial neurons preferring high spatial frequencies (though less than that of non-robust models). Finally, we found that robust models are better models of V1 than non-robust models in terms of their neural response predictivity.

Altogether, although CNNs are some of the best models of the ventral visual stream in terms of neural response predictions, there are still many differences between human and machine perception that need to be improved upon to gain a deeper understanding of our visual system. The results suggest that one way in which our visual system is robust to minute image perturbations is by ignoring (i.e., not encoding) high spatial frequency information in the inputs. They also suggest that explicitly reducing the dimensionality of internal representations (e.g., by penalizing the eigenspectrum so that it decays faster) and reducing the preferred spatial frequency of artificial neurons during task-optimization may improve a model’s adversarial robustness and that this may also lead to better models of V1.

## Results

We performed in-silico electrophysiology experiments and linearly mapped model neurons to macaque V1 neurons. A schematic of the analyses performed in this work is provided in Fig 1. Model layer activations were recorded in response to a set of natural images and a set of Gabor patches in order to obtain the model layer’s eigenspectrum and preferred spatial frequency distribution (Fig 1A and Fig 1B). Using a previously collected set of macaque V1 neural responses [13], we linearly mapped model neurons to V1 neurons (Fig 1C).

**Fig 1.**
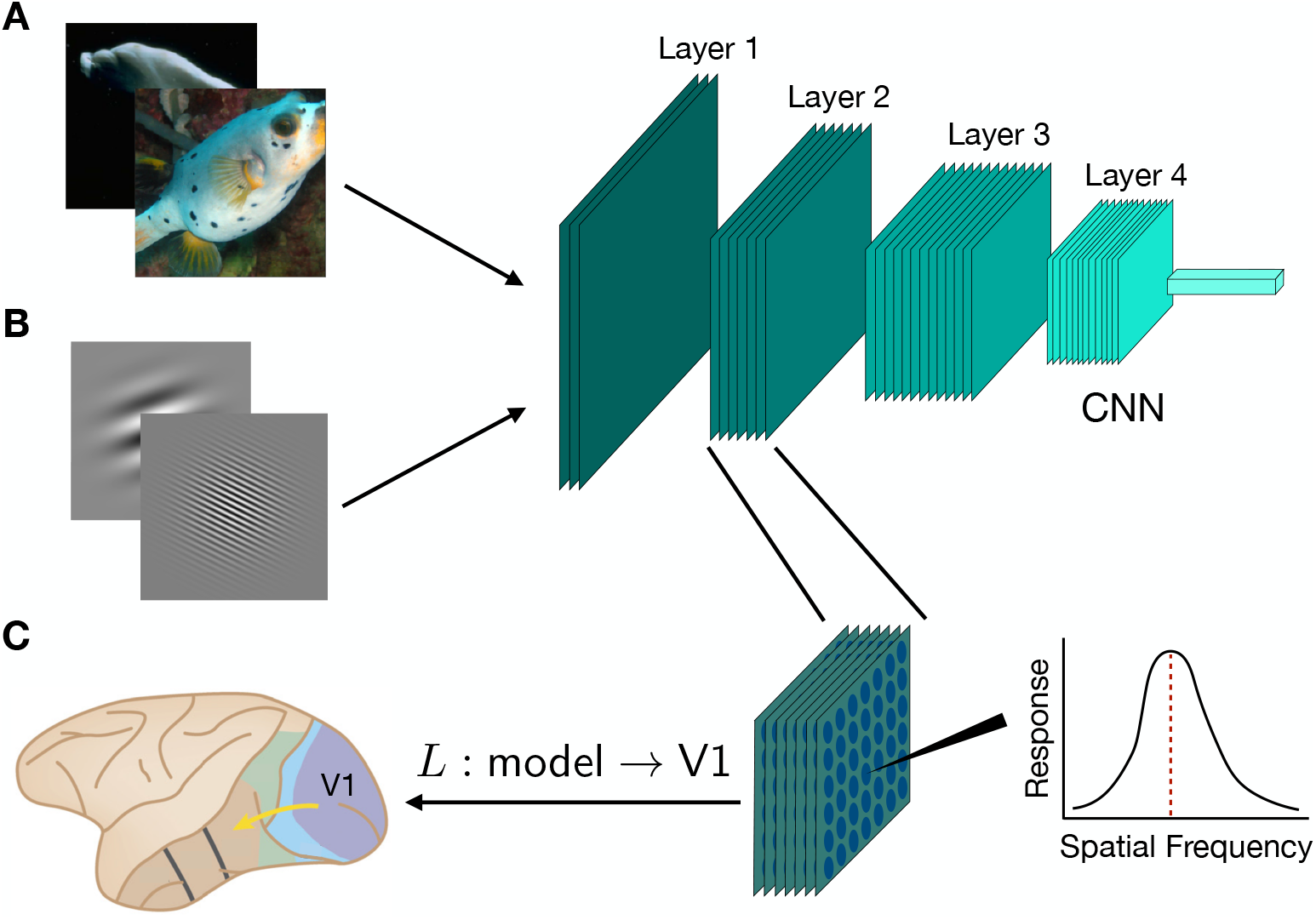
A schematic of the model analyses. **A**. A random set of natural images from the ImageNet database [27] was used to obtain a model’s responses at each layer, from which the eigenspectrum and the power law exponent were computed. **B**. In-silico electrophysiology experiments were performed by presenting models with a set of Gabor patches that varied in spatial frequency, orientation and phase. A spatial frequency tuning curve was then computed using a single artificial neuron’s responses and its preferred spatial frequency is the frequency at which the tuning curve reaches its maximum value. By performing this analysis for each convolutional filter, we computed the distribution of preferred spatial frequencies for a model layer. **C**. A model layer’s responses were linearly mapped (denoted as *L*) to macaque V1 neural responses using partial least squares regression and the linear map’s performance was defined to be the noise-corrected Pearson’s correlation between the model predictions and the observed neural responses. Brain image adapted from [28].

### Eigenspectrum of macaque V1 neural responses also follows a power law with exponent at least one

It was observed by Stringer et al. [26] that the eigenspectrum of mouse V1 neural responses to natural scenes decays according to a power law with exponent close to one. If the power-law-like behaviour of the eigenspectrum is a strong biological constraint, then we would expect that it would generalize across species (e.g., macaques). We therefore computed the eigenspectrum using cross-validated principal components analysis (cvPCA, [26]) of a previously collected set of macaque V1 neural responses to natural scenes [13]. As shown in Fig 2, we found that the eigenspectrum of macaque V1 neural responses follows a power law with exponent close to one, as in mouse V1 [26], suggesting that the power-law-like behaviour of the neural response eigenspectrum does indeed generalize across species.

**Fig 2.**
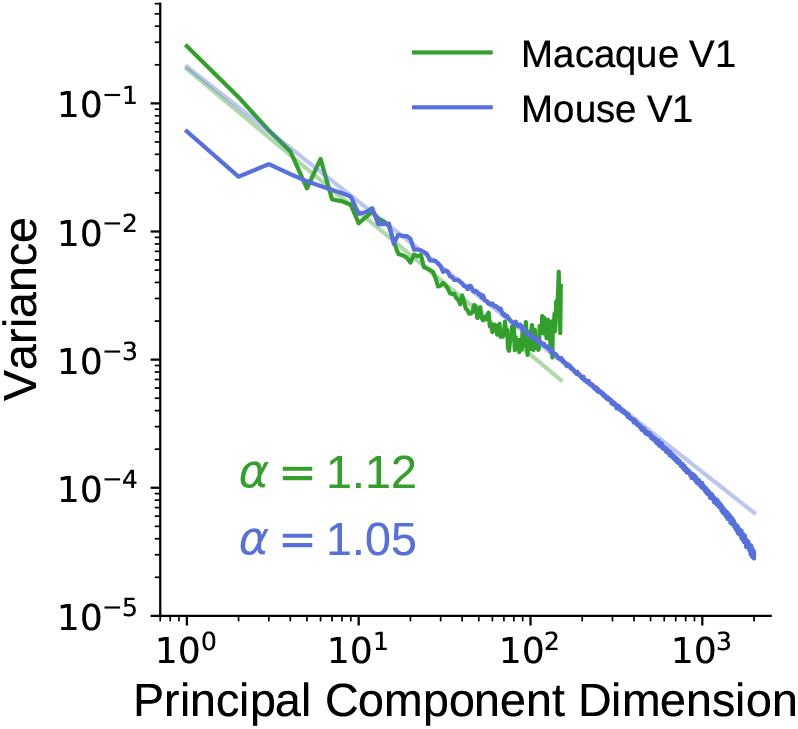
Eigenspectra of macaque and mouse V1 neural responses to natural scenes. Using a previously collected set of macaque V1 neural responses to natural scenes, we computed its eigenspectrum using cvPCA and found that it obeyed a power law and had a power law exponent close to one, similar to the observation in mouse V1 [26]. Light lines are the lines of best fit of each eigenspectrum (in dark colour) in log-log space and the inset indicates the power law exponent, *α*, of each eigenspectrum.

### Eigenspectra of robust models decay slightly faster than those of non-robust models, but slower relative to those observed in neurophysiology

The theory of Stringer et al. [26] predicts that if the eigenspectrum of a system’s neural responses to natural scenes decays with power law exponent less than one (i.e., *α* < 1), then the system will be affected by small perturbations to the inputs, suggesting that this phenomenon is related to the existence of adversarial examples, where human-imperceptible perturbations to an image can cause a model to mis-classify the image even though the unperturbed image could be classified correctly [15, 26]. This implies that the internal representations of non-robust CNNs are greatly affected by “small” image perturbations. Motivated by this theory, we asked whether or not the power law exponents of the eigenspectra of the representations of robust models were higher and closer to one than those of non-robust models.

Here and in the subsequent two sections, we focus on two model architectures - ResNet-50 and ResNet-18 [6]. Both of these task-optimized architectures have been shown to achieve good neural predictivity [11], good task performance [6] and are architectures that have been previously trained with and without robustness penalties. For a particular layer in each model, we computed the eigenspectrum of the artificial neural responses to random sets of approximately 3000 natural images from the ImageNet validation set images [27]. As shown on the left of Fig 3A and Fig 3B, we found that the eigenspectrum of a particular layer of the robust model has a larger power law exponent than that of its non-robust counterpart. In fact, across model layers, we found that the power law exponents of the robust models were higher than those of their non-robust counterparts, as shown on the right of Fig 3A and Fig 3B. These findings are consistent with the theory of Stringer et al. [26], as the models with higher power law exponents are slightly more robust to small image perturbations. We note, however, that the power law exponents of robust models are much less than one and still mismatch with that of V1, consistent with the fact that there still exist image perturbations that can fool these robust models.

**Fig 3.**
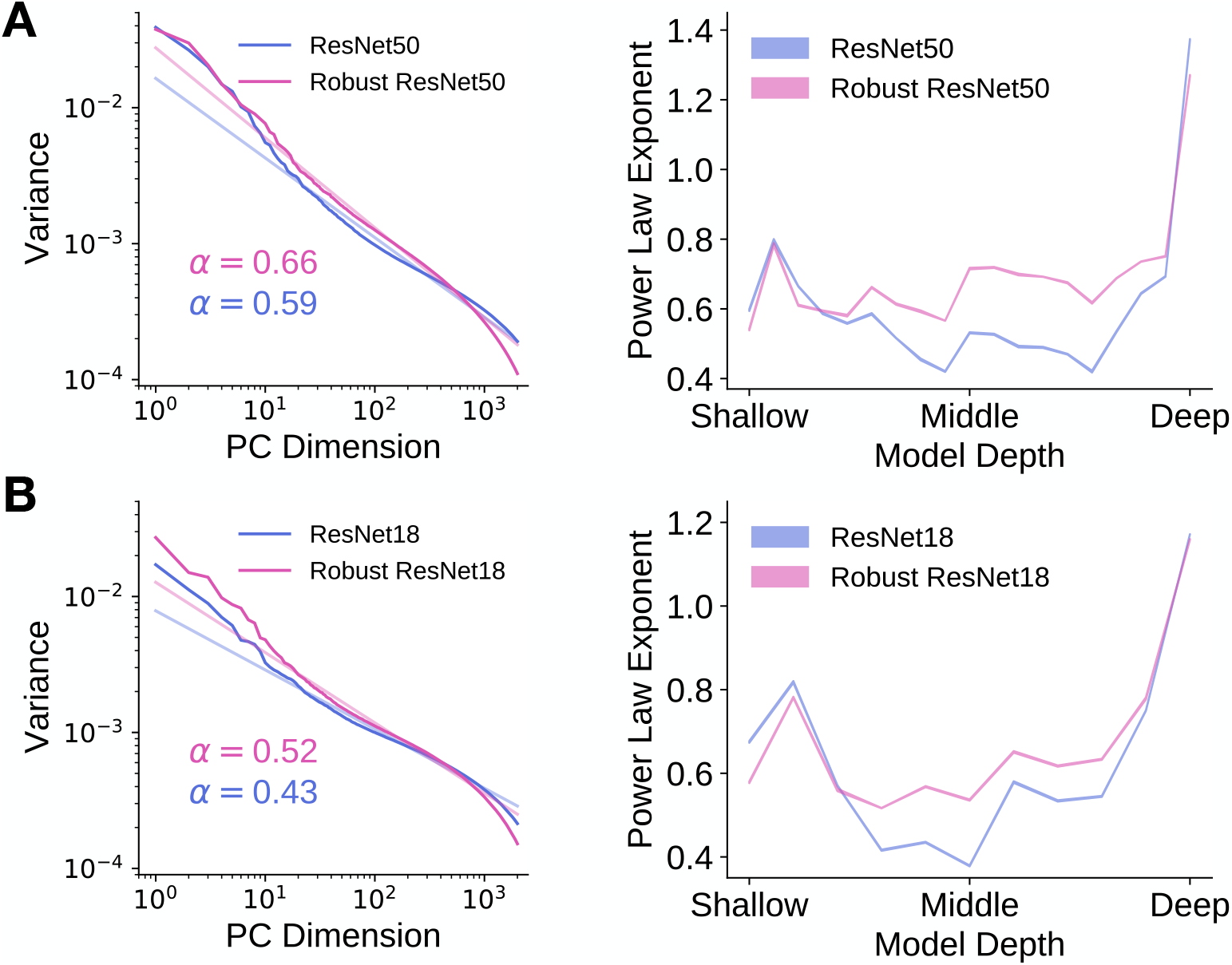
The internal representations of robust models is slightly lower dimensional than those of non-robust models. The eigenspectrum of the robust model decays slightly faster than that of the non-robust model. The eigenspectrum of the artificial neural responses in each layer of a model to a random subset of 2816 ImageNet validation set images was computed using principal components analysis and the power law exponent was computed by obtaining the slope of the line of best fit to the variances within the principal component range of 10 and 1000. **A**. Left: We plot the eigenspectrum for the most “V1-like” layer (as determined by neural predictivity) of a robust and a non-robust ResNet-50. Light lines are the lines of best fit to the eigenspectrum (in dark colour) in log-log space between principal component dimensions 10 and 1000. The inset indicates the estimated power law exponent of the eigenspectrum. Right: Power law exponents of model layers for ResNet-50. Shaded regions (too small to be visible) indicate standard deviation across random ImageNet validation set subsets. **B**. As in **A**, but for ResNet-18.

### Preferred spatial frequency distributions of robust models are more similar to V1 than those of non-robust models

As described above, we observed that the eigenspectra of the internal representations of two robust models decayed slightly faster than those of their non-robust counterparts. Since the eigenspectra for the robust models decay slightly faster than those of the non-robust model, the representations learned by the robust models are lower-dimensional than those learned by the non-robust models. One possible reason for the lower-dimensionality is that “fine stimulus features” may not be contributing to the artificial neural response variance for the large principal component dimensions of robust models as much as they do for the non-robust models.

We therefore hypothesized that non-robust models extract image features that are of higher spatial frequencies than those extracted by robust models. This would mean that its “V1-like” layer (defined as the model layer that has the highest V1 neural response predictivity) consists of a large proportion of artificial neurons tuned to mid to high spatial frequencies. We tested this hypothesis by performing in-silico electrophysiology experiments to estimate the spatial frequency tuning of artificial neurons in each model. We first generated fixed-size Gabor patches of ten orientations, ten spatial frequencies and ten phases, of which a few examples are shown in Fig 4A. The Gabor patches were then presented to the models and artificial neural responses were obtained from the most “V1-like” model layer. Using these responses, we computed spatial frequency tuning curves, of which a few examples are shown in Fig 4B. The preferred spatial frequency of an artificial neuron was then defined to be the spatial frequency at which the tuning curve achieves its maximum value.

**Fig 4.**
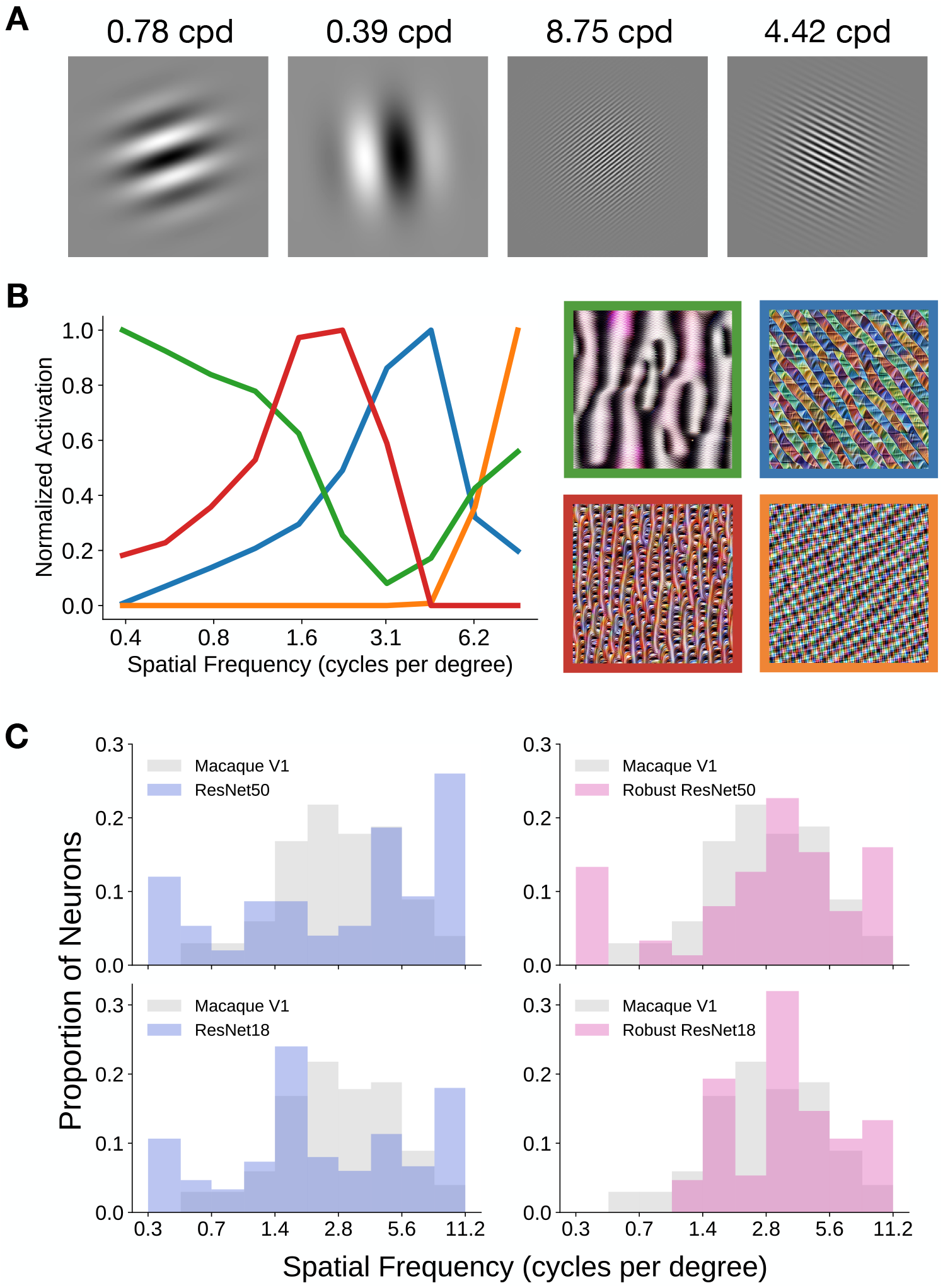
Assessing the spatial frequency tuning of models. We performed in-silico electrophysiology experiments to extract spatial frequency tuning curves of representative artificial neurons from the most “V1-like” layer of a model, as determined by neural predictivity. The field of view of these models was assumed to be 6.4^*°*^ per image. **A**. A few example Gabor patch stimuli of different orientations, phases and frequencies, that were used in the in-silico electrophysiology experiments. cpd: cycles per degree. **B**. A few example spatial frequency tuning curves are plotted, each corresponding to an artificial neuron (i.e., convolutional filter). The images are the stimuli that optimally excite the corresponding channel in the output of the most “V1-like” ResNet-50 model layer. **C**. Preferred spatial frequency distributions for robust and non-robust models from one in-silico experiment. We can see that the distributions of the robust models are more similar to that of macaque V1 cells than the distributions of non-robust models. Top: Robust and non-robust ResNet-50. Bottom: Robust and non-robust ResNet-18.

To gain more intuition into the image features that are extracted by artificial neurons that prefer various spatial frequencies, we generated stimuli that maximally excite channels of the most “V1-like” model layer. We show spatial frequency tuning curves of example artificial neurons and their associated optimal stimuli in Fig 4B. As can be seen, the optimal stimulus for artificial neurons that respond maximally to high spatial frequencies contain high frequency image features. On the opposite end of the spectrum, the optimal stimulus for artificial neurons that respond maximally to low spatial frequencies contain relatively low spatial frequency content.

We constructed the preferred spatial frequency distribution of a model by aggregating the preferred spatial frequencies across the artificial neurons and found that robust models had distributions more similar to that of cells in the foveal area of macaque V1 than those of non-robust models. From the distributions shown in Fig 4C, we observed that both robust and non-robust models have many artificial neurons that are tuned to the highest spatial frequency bin (we assumed the field of view of these models was 6.4 degrees of visual angle, as in the work of Cadena et al. [13]; 56 cycles per image ≈56 cycles */* 6.4 degrees = 8.75 cycles per degree). Comparing these distributions with that of cells in the foveal area of macaque V1 (cf. Fig 6 in De Valois et al. [29]), we note that there are a relatively small number of cells in macaque V1 that are tuned to high spatial frequencies (coloured in gray in Fig 4C), suggesting that this is one way (of many) in which current task-optimized CNN models of the ventral visual stream deviate from the neurophysiology.

Although both robust and non-robust models have preferred spatial frequency distributions that are quite unlike that of macaque V1 foveal cells, the distributions of robust models were more similar to that of macaque V1 than the distributions of non-robust models (Fig 4C). To quantify this similarity, we used a metric based on the maximum absolute difference between the two cumulative distributions (see Methods), where smaller scores indicate that the two distributions are dissimilar and larger scores indicate that the two distributions are similar. For the ResNet-50 architecture, the non-robust model had a score of 0.763± 0.033, whereas the robust model had a score of 0.817 ± 0.032. For the ResNet-18 architecture, the non-robust model had a score of 0.771 ± 0.039, whereas the robust model had a score of 0.790 ± 0.036. The error in all cases denotes the standard deviation across 1000 in-silico electrophysiology experiments.

### Robust models better predict macaque V1 neural responses than non-robust models

We observed that the power law exponent and the preferred spatial frequency distribution of robust models are closer to those of macaque V1, suggesting that robust models better predict V1 neural responses than non-robust models, which we found to be the case. For each model, we assumed its field of view was 6.4 degrees of visual angle, as in prior work [13]. For each model layer, we performed a partial least squares regression to find a linear mapping between the model features and the macaque V1 neural responses, consistent with the procedure described in prior work [8, 11, 30, 31]. The goodness-of-fit of the linear mapping was defined as the correlation between the predicted and the observed neural responses, noise-corrected by the Spearman-Brown corrected cross-trial correlation (i.e., internal consistency) of each neuron. As expected, the feature space provided by the model layers between the shallow and middle portions of the models best corresponded to macaque V1 neural responses, consistent with prior work [13, 14]. Furthermore, as shown in Fig 5, we found that robust models provided feature spaces that better correspond to macaque V1 neural responses than those of non-robust models (*p <* 0.01 for both architectures).

**Fig 5.**
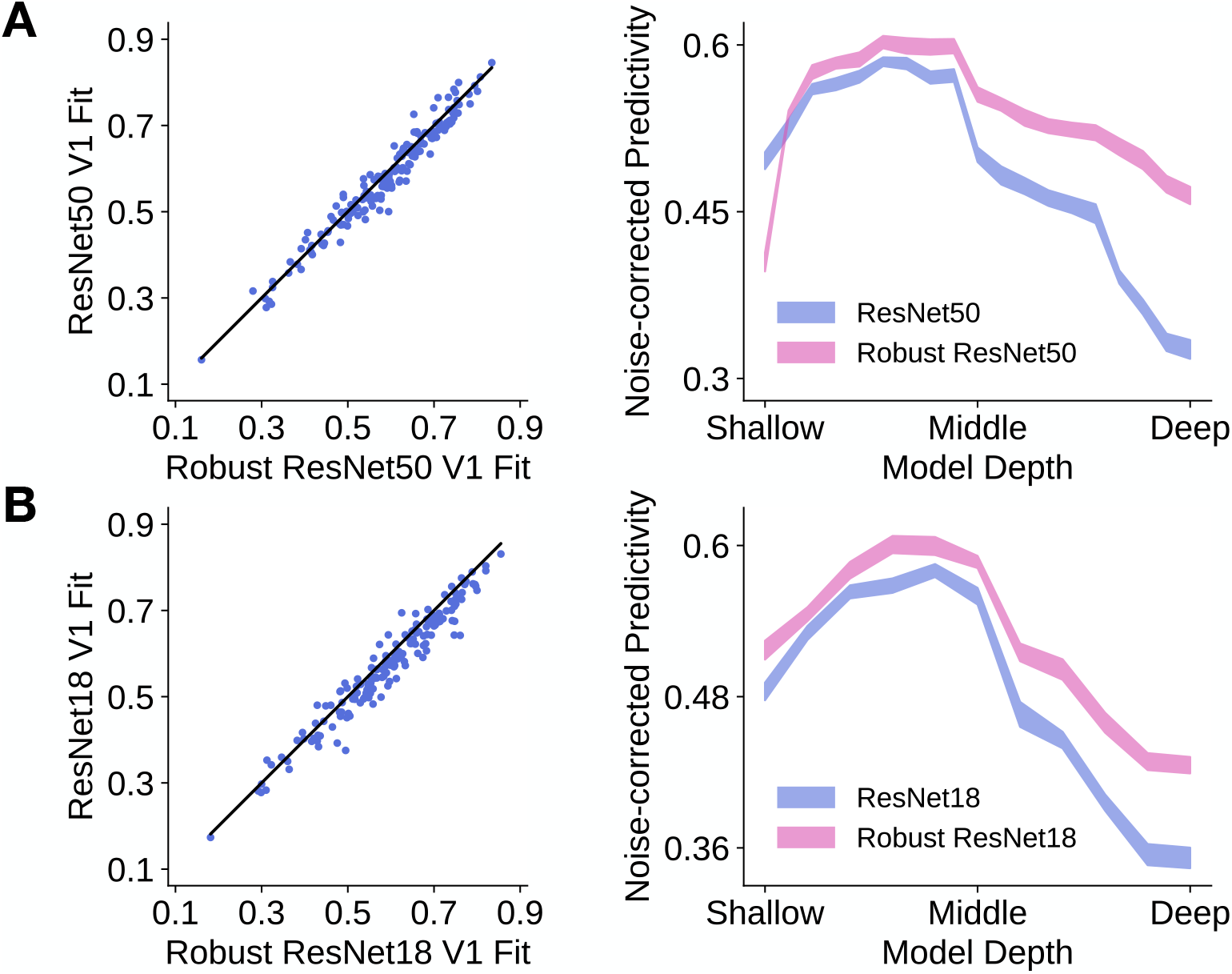
Robust models better predict macaque V1 neural responses than non-robust models. The noise-corrected predictivity for a neuron was defined to be the correlation between the predicted and observed responses, corrected by the neuron’s reliability. **A**. Left: For most neurons (represented by each dot), robust ResNet-50 has higher neural predictivity than non-robust ResNet-50. The black line denotes the identity line. Right: Median neural predictivity across neurons of a robust and non-robust ResNet-50 across model layers. Shaded region indicates the standard deviation of the median neural predictivity across 20 train/test image splits. **B**. As in **A**, but for robust and non-robust ResNet-18. For both ResNet-50 and ResNet-18, the neural predictivities for the robust models are significantly better than those of the non-robust models (*p <* 0.01 for both architectures). Statistical significance was determined by bootstrap resampling of the neurons (with replacement) 10 000 times.

### Adversarial robustness is correlated to V1 predictivity and is not correlated to power law exponent

We observed that two instances of the CNN class of models (ResNet-18 and ResNet-50) that were adversarially trained better corresponded to macaque V1 neural responses than their non-robust counterparts. In addition, the robust models had larger power law exponents across model layers. We next asked whether these observations extended across a wider range of CNN architectures. We therefore performed a large-scale benchmarking of 40 models to ascertain whether or not there was a relationship between a model’s robustness, defined by its accuracy on adversarially perturbed images from the ImageNet validation set [27], and its V1 neural response predictivity. We also compared each model’s robustness with its power law exponent, which was obtained from the model layer that had the highest macaque V1 neural predictivity, determined by partial least squares regression.

Across the set of 40 models, we observed that a model’s adversarial accuracy was strongly correlated to its V1 neural response predictivity (*R* = 0.772, *p <* 0.001). This correlation was robust to the linear regression procedure used (S1 Fig A shows the relationship when ridge regression was used instead). This corroborates prior work of Dapello et al. [30], who had similar observations using a slightly different set of models and a different macaque V1 neural response dataset.

When comparing adversarial accuracy to power law exponent across models, we found a weak relationship between these two quantities (*R* = 0.362, *p* = 0.022). This relationship, however, was not robust to the linear regression procedure used, as shown in S1 Fig B. Although there was not a strong linear relationship *across* models, we found that when comparing a robust model with its non-robust counterpart (i.e., where both models have the same architecture, but one is “robustified” using robustness penalties), the robust model generally had a higher power law exponent. This is shown by the purple lines pointing to the upper right in S1 Fig B. This result is consistent with the predictions of the theory proposed by Stringer et al. [26].

### Higher model robustness is associated with higher alignment with V1 of their preferred spatial frequency tuning distributions

Previously, we described two robust models whose preferred spatial frequency tuning distributions of their most “V1-like” model layers were more like that of macaque V1 than the distributions of non-robust models. In particular, robust models had more artificial neurons that preferred “middle” spatial frequencies (i.e., approximately 3 cpd). We next investigated whether or not this pattern extended across a larger set of architectures. We found that the adversarial accuracy of a model was weakly correlated to its spatial frequency score (*R* = 0.544, *p <* 0.001), indicating that the image features extracted by more robust models have spatial frequencies that might be more aligned with the image features extracted by V1. This result was robust to the linear regression procedure used to select the “V1-like” model layer, as shown in S2 Fig A.

Finally, as shown in Fig 7B, we found that the more similar a model’s preferred frequency distribution is to that of macaque V1, the higher the model’s macaque V1 neural response predictivity (*R* = 0.663, *p <* 0.001). This indicates that our metric for the similarity of a preferred spatial frequency distribution to that of macaque V1 can serve as a reasonable proxy for how good a model is of V1. Of course, this metric can be combined with several other metrics concerning other phenomena of V1, as in work by Marques et al. [31]. This result was also robust to the linear regression procedure used to select the “V1-like” model layer, as shown in S2 Fig B.

**Fig 6.**
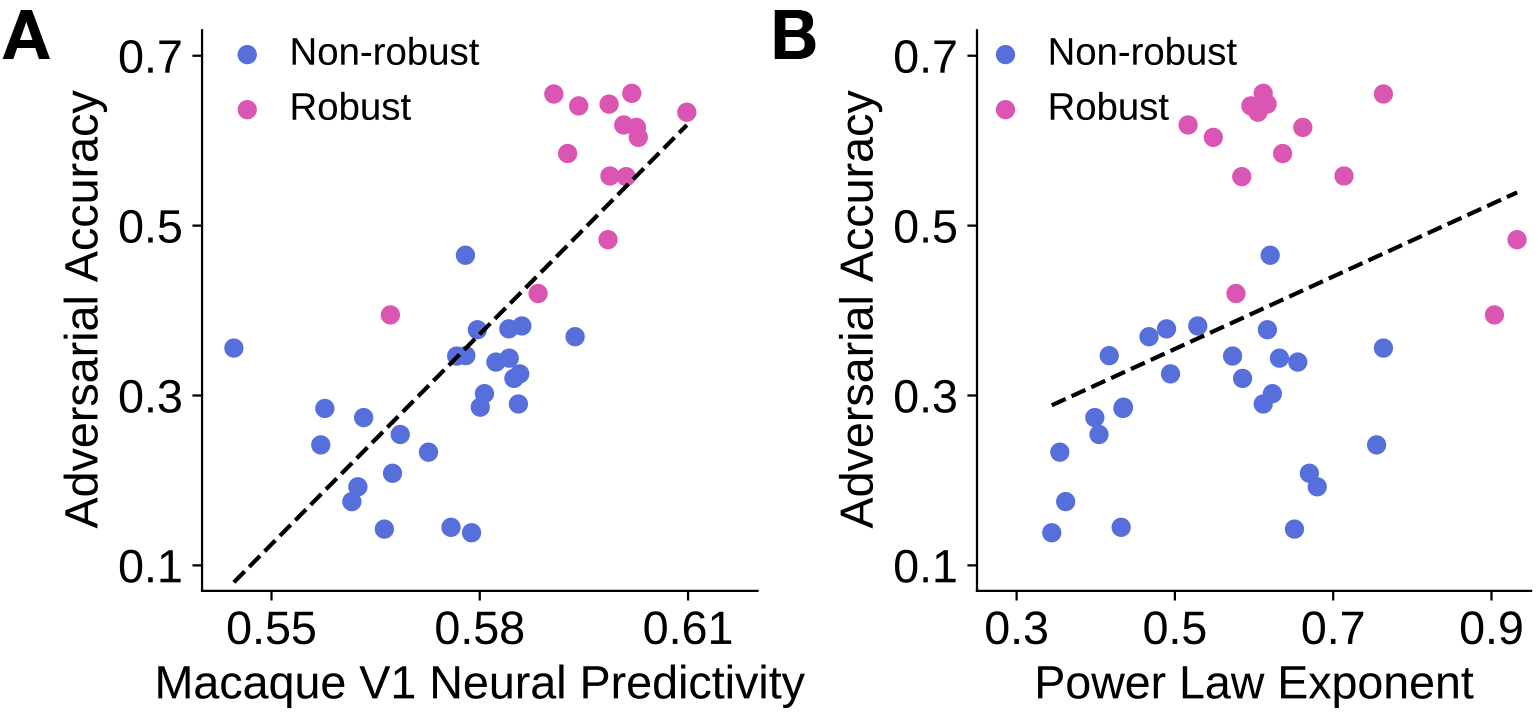
Adversarial accuracy is correlated with V1 neural predictivity and is not correlated with power law exponent. Each model is represented by a dot in each subfigure, with blue denoting non-robust models and pink denoting robust models. Dashed line indicates the line of best fit through the data points. **A**. A model’s adversarial accuracy is plotted against its (maximum) V1 neural response predictivity. As mentioned previously, neural predictivity was defined to be the noise-corrected Pearson correlation between the predicted and observed neural responses. **B**. A model’s adversarial accuracy is plotted against the power law exponent of its most “V1-like” layer, which was determined by neural predictivity.

**Fig 7.**
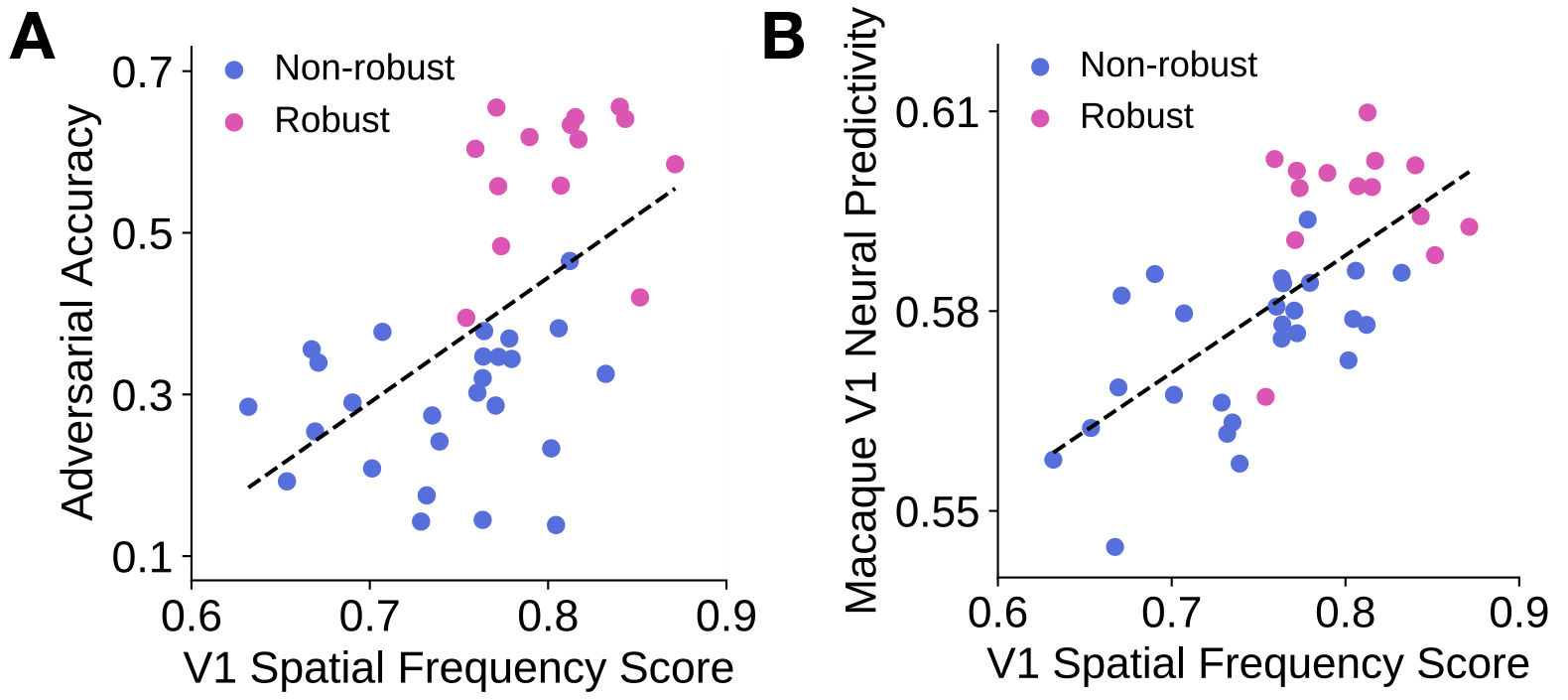
V1 spatial frequency score is somewhat correlated with adversarial accuracy and is correlated with V1 neural predictivity. Each model is represented by a dot in each subfigure, with blue denoting non-robust models and pink denoting robust models. Dashed line indicates the line of best fit through the data points. **A**. A model’s adversarial accuracy is plotted against its V1 spatial frequency score, which denotes the similarity between a model’s preferred spatial frequency distribution and that of macaque V1 cells. **B**. A model’s maximum macaque V1 neural response predictivity is plotted against its V1 spatial frequency score.

**Fig 8.**
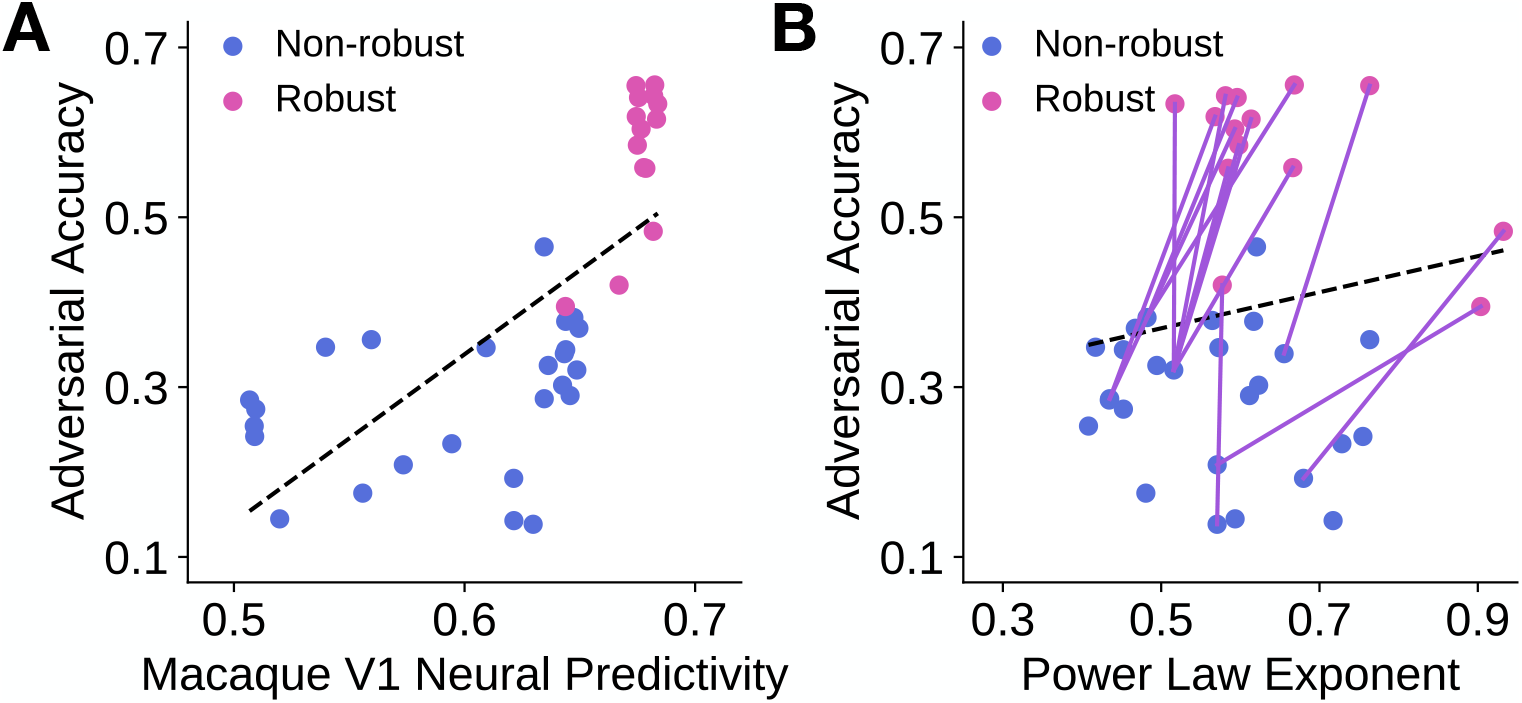
Adversarial accuracy is correlated with V1 neural predictivity obtained via cross-validated ridge regression and is not correlated with power law exponent. Each model is represented by a dot in each subfigure, with blue denoting non-robust models and pink denoting robust models. Dashed line indicates the line of best fit through the data points. **A**. A model’s adversarial accuracy is plotted against its (maximum) V1 neural response predictivity. As mentioned previously, neural predictivity was defined to be the noise-corrected Pearson correlation between the predicted and observed neural responses. **B**. A model’s adversarial accuracy is plotted against the power law exponent of its most “V1-like” layer, which was determined by neural predictivity using ridge regression. Each purple line connects two models of the *same* architecture, where one is trained *without* robustness penalties (blue) and the other is trained *with* robustness penalties (pink).

**Fig 9.**
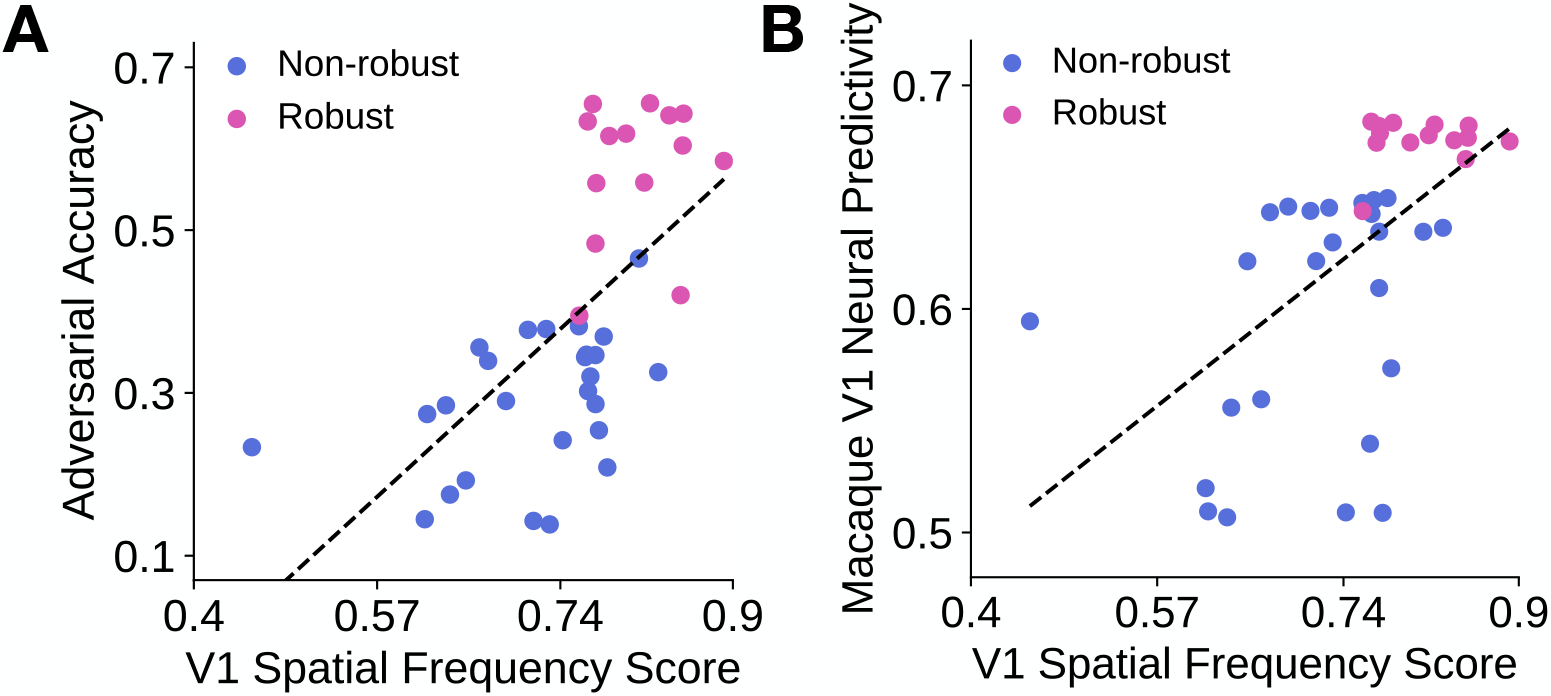
V1 spatial frequency score is correlated with adversarial accuracy and is correlated with V1 neural predictivity obtained via cross-validated ridge regression. Each model is represented by a dot in each subfigure, with blue denoting non-robust models and pink denoting robust models. Dashed line indicates the line of best fit through the data points. **A**. A model’s adversarial accuracy is plotted against its V1 spatial frequency score, which denotes the similarity between a model’s preferred spatial frequency distribution and that of macaque V1 cells. **B**. A model’s maximum macaque V1 neural response predictivity is plotted against its V1 spatial frequency score.

## Discussion

Task-optimized CNNs are susceptible to human-imperceptible adversarial perturbations. In this work, we investigated properties of these CNNs related to robustness to these perturbations in order to gain more insights as to why these models are so brittle. The theory of Stringer et al. [26] suggested that the power law exponent of the eigenspectrum of a set of neural responses (to natural scenes) may be indicative of how prone a system is to small stimulus perturbations, where power law exponents larger than and closer to one would be indicative of a stimulus-neural response mapping that is less susceptible to small input perturbations [26]. We show that the eigenspectra of mouse and macaque V1 neural responses obeyed the theory’s predictions and both followed a power law with exponent at least one. Moreover, we found that models more robust to image perturbations had larger power law exponents than those of non-robust models, consistent with the theory’s predictions. However, they decayed more slowly relative to the neurophysiology. Since a slow decay of the eigenspectrum suggests that substantial model response variance is related to the encoding of fine stimulus features, we performed in-silico electrophysiology experiments in order to assess the spatial frequency tuning of these models and found that models had a large proportion of neurons tuned to high spatial frequencies. Furthermore, robust models had preferred spatial frequency tuning distributions that were more like that of macaque V1 cells and also improved macaque V1 neural response predictions. Taken together, these results describe another way in which machine perception differs from human perception and also suggests that one way in which our visual system achieves robustness to small image perturbations is by ignoring high spatial frequency information in an image.

Our result that a large proportion of artificial neurons are tuned to high spatial frequencies is consistent with other findings, providing additional evidence that there are differences in the image features used by humans and by machines for image classification. For example, Geirhos et al. [22] showed that when image classification-trained models were presented with ambiguous images, they tended to classify images based on the images’ “texture” properties as opposed to their “shape” properties. This is in contrast to humans, who generally classified the ambiguous images based on their “shape” properties. Humans are, by definition, invariant to adversarial perturbations. Ilyas et al. [23] suggested that these perturbations are in fact “non-robust” features—features that are only weakly correlated with an image label, but still provide useful information for the model to learn a good image-label mapping. To show this, the authors constructed a dataset where only non-robust features were useful for the task. As humans are invariant to these non-robust features, the dataset appears to be completely misclassified (cf. Fig 2 in Ilyas et al. [23]). Models that were trained using this “non-robust” dataset attained non-trivial accuracy on a normal test set (i.e., test set images that were not adversarially perturbed), providing evidence that these non-robust features—features that humans are invariant to and presumably do not use—are used by models for image classification.

The model’s preference for high spatial frequencies (relative to that in macaque V1) is also consistent with work that investigated model robustness to image corruptions through the lens of Fourier analysis on images. Yin et al. [25] found that even when models were trained on images that were strongly high-pass filtered, models were able to achieve non-trivial accuracy on ImageNet, indicating the fact that models can detect high-frequency image components that are both useful for the image-classification task and imperceptible to humans (cf. Fig 1 in Yin et al. [25]). They additionally found that adversarially training models and training models with Gaussian data augmentation both resulted in models that were less sensitive to high-frequency noise, but more sensitive to low to mid frequency noise, suggesting that training models in these ways results in a weaker dependence on high-frequency image components.

We observed that models that better corresponded to macaque primary visual cortex also had improved adversarial robustness, suggesting that building more “V1-like” models would improve model robustness. One strategy to develop models that are more “V1-like” is to explicitly optimize the artificial neural representations to be more like the representations obtained from V1, while simultaneously optimizing for task performance. One example of this strategy is work by Li et al. [32], who showed that models can be regularized by optimizing model representations to be similar to those computed using neural responses to natural scenes in primary visual cortex of mice. The authors showed that by incorporating this regularization into the objective function for image classification, model robustness can be improved. Another strategy to improve model robustness could be to build into models known properties of the visual system, as humans are invariant to small image perturbations. For example, Dapello et al. [30] constructed a module based on known properties of primary visual cortex, such as the distributions of preferred orientation and of spatial frequency. The authors then showed that prepending this module to state-of-the-art CNN architectures and optimizing the whole model (except for the module) to perform ImageNet classification can improve adversarial robustness to small image perturbations.

The work described above, however, do not provide normative explanations for the characteristics observed in primary visual cortex (i.e., how these characteristics arose in the first place), as the V1 properties are either learned in a data-driven manner or hard-coded into the models. Our results suggest that such V1 properties as the power law exponent and preferred spatial frequency tuning distribution may arise in order to be robust to high-frequency noise or minute input perturbations. Furthermore, they could provide insight into objective functions or constraints leading to improved robustness and to the phenomena observed in primary visual cortex. Recall our observation that when a model is trained with robust optimization algorithms, the power law exponents in shallow and middle convolutional layers increases and is slightly closer to one (Fig 3). This suggests that it may be useful to explicitly optimize the eigenspectrum of the features in the shallow and the middle layers to have a power law exponent closer to one, while simultaneously optimizing for task performance. Nassar et al. [33] have made progress in this direction. They introduced a novel regularization term that explicitly penalizes eigenspectra which do not have a power law exponent of one and showed that this can improve the adversarial robustness of CNNs trained on a small dataset of handwritten digits. Important future work, we believe, would be to incorporate these regularization methods in a computationally tractable manner in large-scale image classification tasks, as higher performance on such tasks is associated with more quantitatively accurate models of the ventral visual stream [8, 11, 14].

We also observed that more robust models had preferred spatial frequency distributions more aligned with that of primary visual cortex. In particular, there was a larger proportion of artificial neurons in robust models than in non-robust models that preferred spatial frequencies in the middle (Fig 4B and Fig 7A). The development of constraints or regularization methods to tune the preferred frequency distribution is, to our knowledge, an open problem. However, it may be the case that one does not need to explicitly constrain the convolutional filters to prefer particular spatial frequencies. Instead, one could alter the image statistics during training to bias models to learn convolutional filters that extract features across a larger extent of an image. In particular, it is known that infant visual acuity is poor early in development as a result of retinal immaturities and improves over time [34, 35]. This means that early in development, the visual cortex of infants effectively receives images of low spatial resolution as input, which increase in resolution over time. Training CNNs to perform face recognition with blurred inputs has been shown to result in convolutional filters of lower spatial frequencies [36]. We leave it to future work to investigate the implications of a developmental sequence of image resolutions during task-optimization for the preferred spatial frequency distribution of and the adversarial robustness of models.

## Methods

### Macaque V1 neural response dataset

We used a previously collected dataset of neural responses from macaque V1 [13]. We briefly describe the dataset here and refer the reader to the original publication for further details on the experiment [13]. Two macaques were presented with 1450 natural scenes and 5800 synthetically generated images at approximately 2^*°*^ of visual angle. The synthetic images were generated such that their higher-order image statistics (as defined by features in various layers of VGG19) matched those of a natural image. Each stimulus was presented for 60 ms and a linear, 32-channel array was used to record spiking activity. Spike counts were obtained in the 40-100-ms time window post stimulus onset. In our analyses, stimuli that did not have at least two trials per neuron were removed, leaving a total of 6250 stimuli. 166 neurons were obtained for further analyses so that the neural response dataset was of dimensions 6250 stimuli × 166 neurons (after averaging across trials). The Spearman-Brown corrected, cross-trial correlation of each neuron was used in the noise-correction of the predictivity metric for each model layer. We found that the median of these values across neurons was 0.428.

### Convolutional neural network architectures

A set of 26 models trained without robustness penalties and 14 models trained using various robust optimization algorithms [18–21, 37] were used in the analyses. We refer the reader to S1 Table for the complete list of models used in this work. Here we provide more information about the models.

#### Non-robust models

The non-robust models that we used include: AlexNet [2], VGGs [3], ResNets [6], wide ResNets [38], SqueezeNets [39], ShuffleNets [40], DenseNets [41], GoogleNet [42], Inception [43], MobileNet [44] and MNASNets [45]. All of these models were pretrained on ImageNet (in a supervised manner) and accessed through the model zoo of PyTorch [46]. We additionally performed the analyses on a ResNet-50 that was previously trained using an unsupervised algorithm (SimCLR, [47]). This model’s linear evaluation head (which was used for obtaining transfer learning performance on ImageNet) was kept for its robustness evaluation.

#### Robust models

The robust models we used are *somewhat* robust to adversarial perturbations. We consider these models as *somewhat* robust because even after training with these algorithms, there still exists perturbations that can fool these models, evidenced by the fact that they do not achieve the same accuracy on adversarially perturbed images as on unperturbed images (see S1 Table). These models were trained using four different algorithms. We briefly describe the algorithms below.

#### Adversarial training [18, 37]

In adversarial training, one seeks find model parameters that minimize the loss due to the worst-case perturbation:

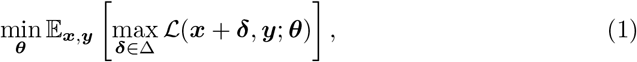

where Δ is the set of “allowed” input perturbations, *δ* is the perturbation, ***ℒ*** is a loss function (e.g., cross-entropy loss), ***x*** is the input (e.g., an image), ***y*** is the output (e.g., image label) and ***θ*** are the model parameters. The inner maximization of Eq (1) is performed using a few steps of projected gradient ascent. More details about this algorithm can be found in S1 Appendix.

Intuitively, this algorithm improves the robustness of models by training models using adversarial examples that are generated during each iteration of training. As additional steps are required to generate adversarial examples, this algorithm can be many times slower than generic training (i.e., supervised training). Adversarially trained models were generously released by Salman et al. [37]. In S1 Table, models trained using this algorithm are denoted as `robust_*`, where * is the name of the base architecture that was adversarially trained.

#### TRADES [19]

This algorithm seeks to improve the adversarial robustness of a model by adding a novel regularization term to the cross-entropy loss. The loss function ***ℒ***_TRADES_ is defined as follows:

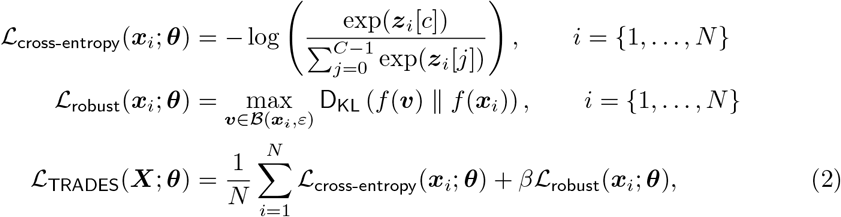

where *N* is the batch size, *C* = 1000 is the number of categories in Image**Net, *z***_*i*_ ∈ ℝ^*C*^ are the model outputs (i.e., logits) for image ***x***_*i*_, *c* ∈ [0, *C* − 1] is the category index of the image (zero-indexed), 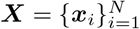 is a batch of *N* images, *𝔅* (***x***_*i*_, *ε*) is a neighbourhood of ***x***_*i*_ of radius *ε, f* (***u***) ∈ ℝ^*C*^ is the vector of log-probabilities for image ***u*** belonging to each of the *C* categories, D_KL_ (·∥·) is the Kullback-Leibler divergence between the two quantities, *β* is the regularization coefficient and ***θ*** are the model parameters.

As in the work of Zhang et al. [19], the maximization in ℒ_robust_ was performed using a few iterations of projected gradient ascent. During each iteration, a new image ***v*** = ***x*** + ***δ*** is computed so as to increase the value of D_KL_ (*f* (***v***) ∥ *f* (***x***)). The steps are nearly identical to those used for the inner maximization in Eq (1). In particular, ℒ(***v***^(*t*)^; ***θ***) := D_KL_ *f* (***v***^(*t*)^) 1 ∥*f* (***x***), and the Project(·, ·) function is that defined for *l*_*∞*_-norm constraints (i.e., perturbations are clipped to be within [− *ε, ε*]).

We trained a ResNet-50 architecture on ImageNet using the loss function defined in Eq (2). The loss function was minimized using stochastic gradient descent (SGD, [48]) with momentum for 80 epochs. We used a batch size of 128, momentum of 0.9 and an initial learning rate of 0.1, which was decayed by a factor of 10 at epochs 25, 45 and 65. The maximization in ℒ_robust_ of Eq (2) was performed using three gradient ascent steps, with a step size of *η* = 4*/*255 × 2*/*3 and the regularization coefficient was set to *β* = 2. The maximum perturbation size allowed was ∥***δ∥*** _*∞*_ ≤4*/*255. Finally, the weight decay was set to 0.0001. The model trained with this algorithm is denoted as `trades_robust_resnet50_linf_4` in S1 Table.

#### Input gradient regularization [21]

This algorithm seeks to improve the adversarial robustness of models by adding a regularization term to the cross-entropy loss. At a high-level, the regularization term penalizes the gradient of the loss function with respect to the input. Concretely, the loss function L_IGR_ is defined as follows:

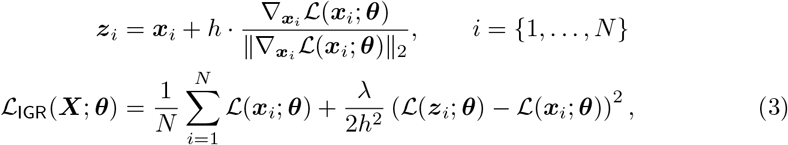

**where ℒ(·; *θ***) is the cross-entropy loss, *λ* = 0.3 is the regularization coefficient, *h* = 0.01 is the finite difference hyperparameter, 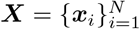 is a batch of *N* images and ***θ*** are the model parameters.

A model with the ResNet-50 architecture was trained to minimize Eq (3) using SGD with momentum for 100 epochs. We used a batch size of 128, momentum of 0.9 and an initial learning rate of 0.1, which was decayed by a factor of 10 at epochs 35, 70 and 90. Finally, the weight decay was set to 0.0001. The model trained with this algorithm is denoted as `igr_robust_resnet50` in S1 Table.

#### Adversarial training for free [20]

As mentioned previously, adversarial examples are generated on each iteration of adversarial training. Thus, assuming that *K* steps are used to generate the adversarial examples during each iteration, adversarial training can be *K* + 1 times slower than generic training. “Free adversarial training” was developed in order to reduce the total time required to train these models. At a high level, gradients with respect to the input are accumulated over multiple training steps circumventing the need to compute the gradient multiple times during each training step.

A model with the ResNet-50 architecture was trained using this algorithm for 23 epochs, where each batch of 256 images was repeated four times during each epoch. SGD with momentum of 0.9 was used and the weight decay was set to 0.0001. The initial learning rate of 0.1 was decayed by a factor of 10 every 30 epochs. The maximum perturbation size allowed was ∥*δ*∥_*∞*_ = 4*/*255 and the step size was also 4*/*255. The model trained with this algorithm is denoted as free robust resnet50 linf 4 in S1 Table.

### Computing power law exponents

#### Power law exponent of macaque V1 neural responses

In order to compute the eigenspectrum of the macaque V1 neural responses to natural scenes, we used cross-validated principal components analysis (cvPCA). It was developed by Stringer et al. [26] to compute the eigenspectrum of neural responses from mouse V1. This algorithm computes unbiased estimates of the eigenvalues (and hence the eigenspectrum) of the population neural responses. Briefly, the algorithm operates as follows:

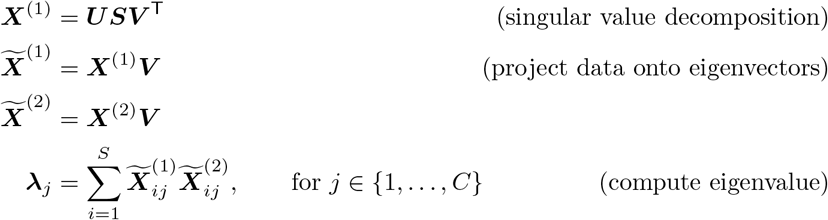

where *S* is the number of stimuli, *N* is the number of neurons, ***X***^(1)^, ***X***^(2)^ ∈ℝ^*S×N*^ are the neural responses for the first and second half of the trials (and averaged across trials), ***V*** ∈ ℝ^*N ×C*^ are the *C* eigenvectors of the covariance matrix of ***X***^(1)^ and ***λ*** ∈ ℝ^*C*^ are the cross-validated eigenvalues associated with each of the eigenvectors (***λ***_*j*_ is the *j*th eigenvalue).

The first step of the cvPCA algorithm computes the eigenvectors of the neural response covariance from one set of the trials. The second and third steps project the neural responses from each half of the trials onto each eigenvector. The final step computes the (scaled) variance of the neural responses when projected onto an eigenvector (that was computed using one half of the trials). Thus, each cross-validated eigenvalue is related to the amount of stimulus-related variance of the neural responses along the eigenvalue’s corresponding eigenvector. The power law exponent was then determined as the negative slope of the line of best fit of the eigenspectrum in log-log space, similar to the procedure described by Stringer et al. [26]. We refer the reader to the original publication for a more detailed mathematical analysis of this method [26]. We ran this algorithm 20 times for the macaque V1 neural response dataset and averaged the eigenvalues computed from each of the 20 runs.

#### Power law exponents of artificial neural responses

The eigenspectrum of artificial neural responses to three random sets of 2816 images from the ImageNet validation set was computed for each model layer. Each image was first resized so that its shorter dimension was 256 pixels and then center-cropped to 224 ×224 pixels.

Images were additionally preprocessed by normalizing each image channel (RGB channels) using the mean and standard deviation that was used during model training. Using these preprocessed images, we extracted activations from several layers of each CNN and computed their eigenspectra using principal components analysis (PCA). We did not use cvPCA, as we did for the macaque V1 neural responses, because artificial neural responses are deterministic. Similar to the procedure described by Stringer et al. [26], the power law exponent was estimated as the negative slope of the line of best fit of the eigenspectrum in log-log space over the principal component indices in the range of 10 to 1000. For the analysis pertaining to the comparison of a model’s robustness with its power law exponent, we summarized each model by the power law exponent of the model layer that best predicted the macaque V1 neural responses.

### Spatial frequency tuning

#### Preferred spatial frequency tuning distribution of models

In order to assess the spatial frequency tuning of artificial neurons, we performed in-silico electrophysiology experiments. Specifically, we first generated Gabor patches of various orientations, spatial frequencies and phases according to the following equation:

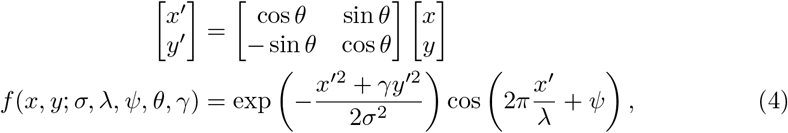

where *σ* = 35 determines the standard deviation of the Gaussian envelope in pixels, *γ* = 1 is the aspect ratio of the Gabor patch, *λ* is the number of pixels for one cycle of the sinusoid, *ψ* is the phase of the sinusoid in radians and *θ* is the orientation of the Gabor patch in radians. We generated Gabor patches with different orientations, phases and spatial frequencies using the following parameters: 10 orientations were evenly spaced between 0^*°*^ and 172.5^*°*^ and 10 phases were evenly spaced between 0^*°*^ and 360^*°*^. The following spatial frequencies were used (in units of cycles per image): 2.5, 3.5, 5, 7.1, 10, 14.1, 20, 28.3, 40, 56. This resulted in 10 × 10 ×10 = 1000 Gabor patch stimuli. Prior to presenting the Gabor patch stimuli to the models, each stimulus was preprocessed by normalizing the RGB channels using the mean and standard deviation that the model was trained on.

Here we describe the method by which we obtained spatial frequency tuning curves of artificial neurons from model layers. The output of a convolutional layer is a matrix of dimensions *C* × *H* × *W*, where *C* is the number of channels (i.e., convolutional filters), and *H* and *W* are the height and width. Since the model layer is convolutional, each artificial neuron in each channel would have the same “tuning”. Intuitively, each artificial neuron in the same channel would detect the same “features” as they are each associated with the same convolutional filter. As a result, one does not need to obtain the tuning for *all* artificial neurons in the output of a convolutional layer. Thus, for a particular channel (i.e., convolutional filter), we computed the tuning for the artificial neuron at the center of the activations matrix, which we denote as the “representative neuron” (i.e., the neuron at location (⌊*H/*2⌋, *⌊W/*2⌋)). The receptive field of the central artificial neuron would cover the Gabor patch stimuli, as the Gabor patch is placed at the center of the model’s visual field.

In order to compute the value of the tuning curve for an artificial neuron at a particular spatial frequency, we averaged the activations of the artificial neuron to Gabor patch stimuli of all orientations and phases with that particular spatial frequency. This was performed for each of the desired spatial frequencies. Once the tuning curves were computed, artificial neurons were further sub-selected according to their tuning curves’ peak-to-peak value. Artificial neurons were kept only if the peak-to-peak value of their tuning curves were greater than zero. The neuron’s preferred spatial frequency was defined to be the frequency at which the tuning curve achieves its maximum value.

To mimic one electrophysiology experiment, we randomly sampled 150 representative neurons (with replacement) from the model layer’s output and obtained each neuron’s preferred spatial frequency, resulting in the model layer’s preferred spatial frequency distribution. We performed 1000 in-silico electrophysiology experiments and therefore obtained 1000 preferred spatial frequency distributions for a particular model layer. Each of these distributions was then compared with that of macaque V1 cells and a score was computed for each distribution, resulting in 1000 scores (see below for the definition of the scoring function). Each model’s score was then defined to be the average score across the 1000 in-silico experiments (each of which is presented in Fig 7A) and the error was defined to be the standard deviation of the scores across the in-silico experiments.

#### Preferred spatial frequency tuning distribution of macaque V1

Using the online tool called “WebPlotDigitizer”, we extracted data from Fig 6 of De Valois et al. [29], which shows the preferred spatial frequency distribution of neurons in the foveal area of macaque V1. The extracted spatial frequency bins were as follows (in units of cycles per degree): 0.35, 0.5, 0.7, 1.0, 1.4, 2.0, 2.8, 4.0, 5.6, 8.0 (with 11.2 as the rightmost bin edge). The extracted cell counts for each spatial frequency bin were as follows: 0, 3, 3, 6, 17, 22, 18, 19, 9, 4.

#### V1 spatial frequency score

Here we describe the metric that we used to assess the similarity between a model’s preferred spatial frequency distribution and that of macaque V1 cells. The score was defined to be the one minus the maximum absolute difference between the two empirical cumulative distributions (similar to the Kolmogorov-Smirnov distance):

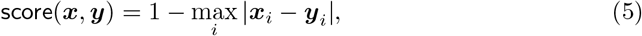

where ***x*** and ***y*** are the empirical cumulative distributions of two different samples and ***x***_*i*_ is the value of the cumulative distribution at the *i*th bin (and correspondingly for ***y****i*). In our case, ***y*** would be the cumulative distribution for the preferred spatial frequency histogram of macaque V1 cells (obtained from the histogram in Fig 6 of De Valois et al. [29]) and ***x*** would be the model’s cumulative distribution for its preferred spatial frequency histogram.

### V1 neural response predictivity

In line with Cadena et al. [13], we first cropped each stimulus to the central 80 pixels and then resized the images to 40× 40 pixels, as we also assumed that each models’ field of view is 6.4 degrees. Each stimulus was then zero-padded up to the image size on which each model was trained. For example, the 40× 40 pixels image would be zero-padded up to 224 × 224 pixels for most models. Each image channel was additionally normalized according to the mean and standard deviation used during model training. Model features were then extracted in response to each preprocessed stimulus.

In order for neural response prediction for each of the 40 models (and their representative model layers) to be more computationally tractable, we first projected the features of each model layer into a 1000-dimensional space using PCA prior to linear fitting. If the number of features was less than 1000 (e.g., there are 512 features in the average-pooling layer of a ResNet-18), the number of principal components used was equal to the number of features. ImageNet validation set images were used to compute the principal components so that the transformation was held constant across all 20 train-test splits during neural fitting [11]. These lower-dimensional stimulus features were then used as input to a partial least squares (PLS) regression procedure with 25 components, consistent with prior work [8, 11, 30, 31]. Each train-test split was generated by randomly selecting 75% of the stimuli to be in the train set and 25% of the stimuli to be in the test set. For the data shown in S1 Fig and S2 Fig, we performed cross-validated ridge regression, where five-fold cross validation (using the train set) was used to obtain the best regularization coefficient from {0.01, 0.1, 1, 10}.

The noise-corrected predictivity metric for a neuron was defined to be the Pearson’s correlation between the neuron’s response predictions and the observed neural responses divided by the square-root of the Spearman-Brown corrected cross-trial correlation of the neuron’s responses, consistent with prior work [14, 49].

### Model robustness

We defined the robustness of a model to be its classification accuracy on the 50 000 ImageNet validation set images that have been perturbed using a set of white-box adversarial attacks and averaged across the set of adversarial attacks. This is referred to as the model’s “adversarial accuracy”. Adversarial perturbations were generated using projected gradient ascent (PGD, [18]). The algorithm used to generate adversarial images is the same as that used for the inner maximization in Eq (1) of adversarial training. For more details, we refer the reader to S1 Appendix.

If the image perturbations can be as large as possible, one can easily distort the image so that it becomes completely unrecognizable by a human (e.g., by modifying the image so that it looks like noise). Thus, the sizes (i.e., norm) of the perturbations, ***δ***, were bounded so that they are imperceptible to humans. In order to generate more variability in the adversarial accuracy across models, we used relatively small adversarial perturbations. Larger perturbations would reduce the adversarial accuracy of most CNNs to chance level. Specifically, the maximum sizes of the perturbations were defined as follows: ∥***δ∥*** _*∞*_ ≤1*/*1020, ∥***δ∥*** _2_ ≤0.15, ∥***δ∥*** _1_ ≤40. Adversarial examples were generated using projected gradient ascent for 20 steps. The step size was set to be *ε* × 2*/*20, where *ε* is the maximum allowed size of the perturbation. This white-box adversarial attack method and these perturbation constraints were also used in previous work that compared adversarial accuracy with V1 neural response predictivity [30]. We used a Python package known as Foolbox [50] to evaluate the robustness of the CNNs.

### Optimal stimulus visualization

As in prior work, we optimized the discrete Fourier transform (DFT) of the input to maximize the activations of a particular channel of a CNN layer. Concretely, we maximized the softmax of the average activation in the desired channel of the output of a convolutional layer:

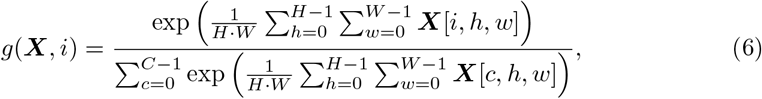

where *i* is the index of the channel we would like to maximize the activations of, *H* and *W* are the height and width of the outputs of the convolutional layer, *C* is the total number of channels in the output of the convolutional layer and ***X***∈ ℝ^*C×H×W*^ is the output of the convolutional layer. Eq (6) was maximized using the Adam optimizer with a learning rate of 0.05. We used a Python package known as Lucent (a PyTorch adapation of Lucid [51]) to generate the optimal stimuli.

## Supporting information

**S1 Fig. Relationships between adversarial accuracy, V1 neural predictivity and power law exponent**. In the main text, we showed results where the “V1-like” layer of a model was obtained via partial least squares regression, where we found that adversarial accuracy correlated with V1 neural predictivity and weakly correlated with power law exponent (recall that the power law exponent for a model was obtained from the model layer that best predicted the macaque V1 neural responses). Here, we show the same figures, but with results obtained via cross-validated ridge regression (where five-fold cross-validation was used to obtain the optimal regularization coefficient). Consistent with our partial least squares regression finding, adversarial accuracy was correlated with V1 neural predictivity (*R* = 0.696, *p <* 0.001). However, when using ridge regression to determine the most “V1-like” model layer, adversarial accuracy was found *not* to correlate with power law exponent (*R* = 0.159, *p* = 0.327). Although there was no linear relationship between adversarial accuracy and power law exponent, we noticed that when comparing two models obtained by training a single architecture with and without robustness penalties, the robust model had higher power law exponents (as shown by the purple lines pointing to the upper right in the figure, indicating higher adversarial accuracy and higher power law exponent). This is consistent with the theory of Stringer et al. [26].

**S2 Fig. Relationships between adversarial accuracy, V1 spatial frequency score and V1 neural predictivity**. In the main text, we showed the relationship between a model’s adversarial accuracy, its V1 spatial frequency score and its maximum V1 neural predictivity when partial least squares regression was used to obtain a model’s “V1-like” layer. Here, we show results pertaining to these relationships, obtained via cross-validated ridge regression. Qualitatively, the results are the same as those described in the main text. Here, we find that a model’s adversarial accuracy and its V1 spatial frequency score was correlated (*R* = 0.624, *p <* 0.001). Furthermore, a model’s maximum V1 neural predictivity was correlated to its V1 spatial frequency score (*R* = 0.565, *p <* 0.001).

**S1 Table. Model performance on ImageNet and V1 neural predictivity**. Table 1 lists all the models we used, their macaque V1 neural response predictivity and their top-1 accuracies on ImageNet validation set images which have not been perturbed (i.e., for a perturbation, ***δ***, and for any norm, ∥***δ***∥ = 0) or were adversarially perturbed with different norm constraints on the perturbations: ∥***δ***∥_*∞*_ ≤ 1*/*1020, ∥***δ***∥_2_ ≤ 0.15, ∥***δ***∥_1_ ≤ 40.

**Table 1.**
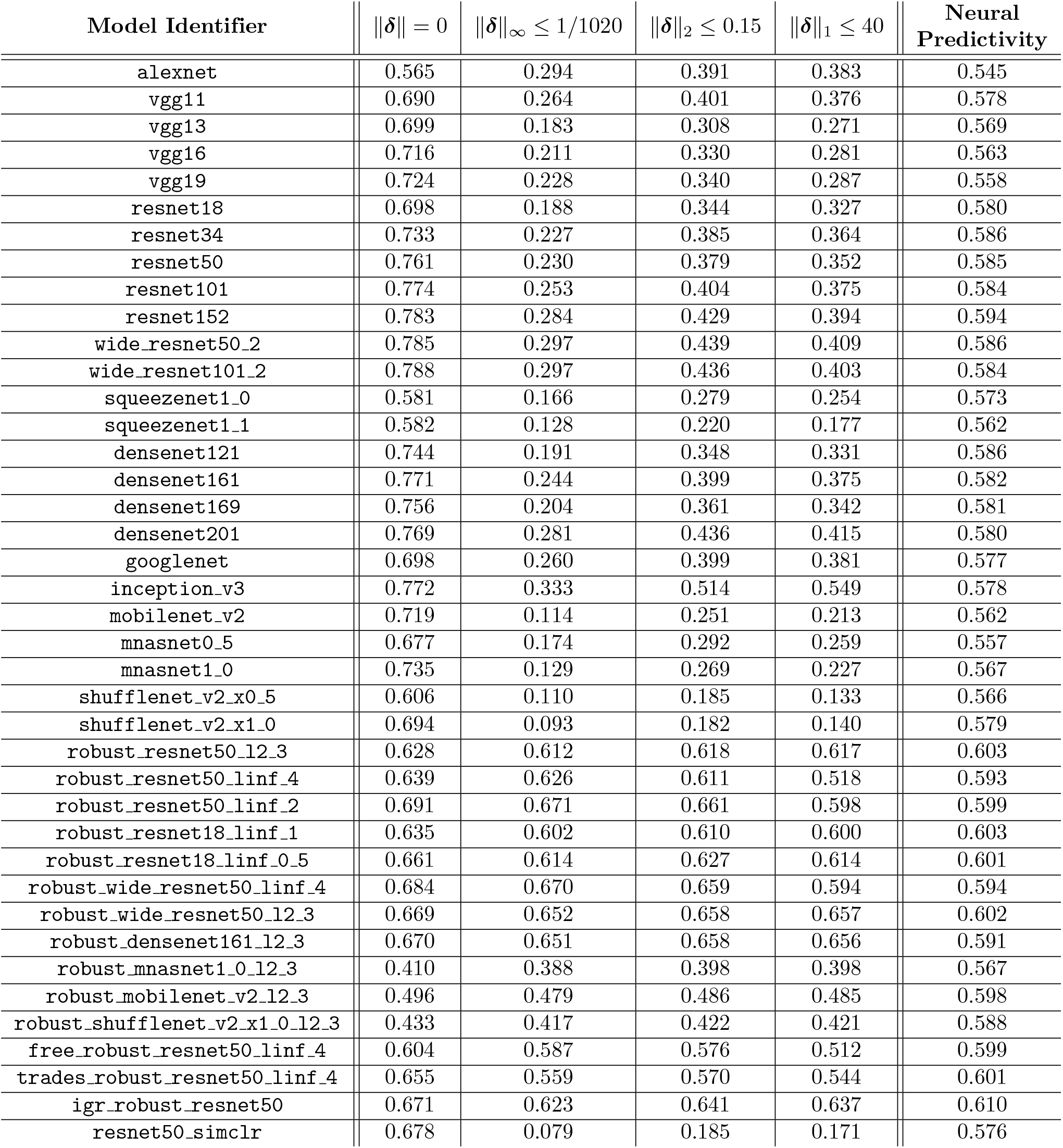
Model neural predictivity (PLS regression) and top-1 accuracies on ImageNet validation set images that have been adversarially perturbed. Each column corresponds to the constraint on the perturbation size. ∥***δ***∥ = 0 corresponds to unperturbed (i.e., “clean”) images. For the models trained to be adversarially robust, the suffix corresponds to the norm constraint imposed on the size of the perturbation during model training. For example, `robust_resnet50_l2_3` corresponds to a ResNet-50 adversarially trained to be robust to perturbations, ***δ***, of size at most **∥ *δ* ∥**_2_ 3 ≤ [37], `igr_robust_resnet50` corresponds to a ResNet-50 trained with input gradient regularization (IGR, [21]) and `resnet50_simclr` corresponds to a ResNet-50 trained with the SimCLR unsupervised loss function [47].

| Model Identifier | $\ \delta\ = 0$ | $\ \delta\ _\infty \leq 1/1020$ | $\ \delta\ _2 \leq 0.15$ | $\ \delta\ _1 \leq 40$ | Neural Predictivity |
| --- | --- | --- | --- | --- | --- |
| alexnet | 0.565 | 0.294 | 0.391 | 0.383 | 0.545 |
| vgg11 | 0.690 | 0.264 | 0.401 | 0.376 | 0.578 |
| vgg13 | 0.699 | 0.183 | 0.308 | 0.271 | 0.569 |
| vgg16 | 0.716 | 0.211 | 0.330 | 0.281 | 0.563 |
| vgg19 | 0.724 | 0.228 | 0.340 | 0.287 | 0.558 |
| resnet18 | 0.698 | 0.188 | 0.344 | 0.327 | 0.580 |
| resnet34 | 0.733 | 0.227 | 0.385 | 0.364 | 0.586 |
| resnet50 | 0.761 | 0.230 | 0.379 | 0.352 | 0.585 |
| resnet101 | 0.774 | 0.253 | 0.404 | 0.375 | 0.584 |
| resnet152 | 0.783 | 0.284 | 0.429 | 0.394 | 0.594 |
| wide_resnet50_2 | 0.785 | 0.297 | 0.439 | 0.409 | 0.586 |
| wide_resnet101_2 | 0.788 | 0.297 | 0.436 | 0.403 | 0.584 |
| squeezenet1.0 | 0.581 | 0.166 | 0.279 | 0.254 | 0.573 |
| squeezenet1.1 | 0.582 | 0.128 | 0.220 | 0.177 | 0.562 |
| densenet121 | 0.744 | 0.191 | 0.348 | 0.331 | 0.586 |
| densenet161 | 0.771 | 0.244 | 0.399 | 0.375 | 0.582 |
| densenet169 | 0.756 | 0.204 | 0.361 | 0.342 | 0.581 |
| densenet201 | 0.769 | 0.281 | 0.436 | 0.415 | 0.580 |
| googlenet | 0.698 | 0.260 | 0.399 | 0.381 | 0.577 |
| inception_v3 | 0.772 | 0.333 | 0.514 | 0.549 | 0.578 |
| mobilenet_v2 | 0.719 | 0.114 | 0.251 | 0.213 | 0.562 |
| mnasnet0.5 | 0.677 | 0.174 | 0.292 | 0.259 | 0.557 |
| mnasnet1.0 | 0.735 | 0.129 | 0.269 | 0.227 | 0.567 |
| shufflenet_v2_x0.5 | 0.606 | 0.110 | 0.185 | 0.133 | 0.566 |
| shufflenet_v2_x1.0 | 0.694 | 0.093 | 0.182 | 0.140 | 0.579 |
| robust_resnet50_l2_3 | 0.628 | 0.612 | 0.618 | 0.617 | 0.603 |
| robust_resnet50_linf_4 | 0.639 | 0.626 | 0.611 | 0.518 | 0.593 |
| robust_resnet50_linf_2 | 0.691 | 0.671 | 0.661 | 0.598 | 0.599 |
| robust_resnet18_linf_1 | 0.635 | 0.602 | 0.610 | 0.600 | 0.603 |
| robust_resnet18_linf_0.5 | 0.661 | 0.614 | 0.627 | 0.614 | 0.601 |
| robust_wide_resnet50_linf_4 | 0.684 | 0.670 | 0.659 | 0.594 | 0.594 |
| robust_wide_resnet50_l2_3 | 0.669 | 0.652 | 0.658 | 0.657 | 0.602 |
| robust_densenet161_l2_3 | 0.670 | 0.651 | 0.658 | 0.656 | 0.591 |
| robust_mnasnet1.0_l2_3 | 0.410 | 0.388 | 0.398 | 0.398 | 0.567 |
| robust_mobilenet_v2_l2_3 | 0.496 | 0.479 | 0.486 | 0.485 | 0.598 |
| robust_shufflenet_v2_x1.0_l2_3 | 0.433 | 0.417 | 0.422 | 0.421 | 0.588 |
| free_robust_resnet50_linf_4 | 0.604 | 0.587 | 0.576 | 0.512 | 0.599 |
| trades_robust_resnet50_linf_4 | 0.655 | 0.559 | 0.570 | 0.544 | 0.601 |
| igr_robust_resnet50 | 0.671 | 0.623 | 0.641 | 0.637 | 0.610 |
| resnet50_simclr | 0.678 | 0.079 | 0.185 | 0.171 | 0.576 |

### S1 Appendix. Additional details on adversarial training

As in the work of Madry et al. [18], the steps to generate adversarial examples from the original image, ***x***, are as follows:

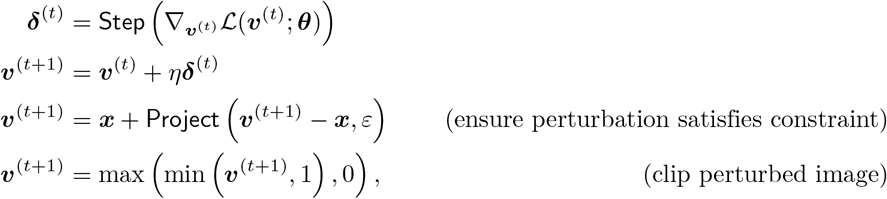

where ℒ (·; ***θ***) is the cross-entropy loss, *η* is the gradient ascent step size, ***v***^(*t*)^ is the perturbed image at step *t* of projected gradient ascent and is initialized to the original image (i.e., ***v***^(0)^ = ***x***) and ***θ*** are the model parameters. Depending on the norm constraint on the perturbations, `Step`(·) and `Project`(·, ·) would be implemented differently.

For *l*_*∞*_-norm constraints, `Step`(·) computes the sign of the gradient and `Project`(·, ·) clamps the perturbation to be within [−*ε, ε*]:

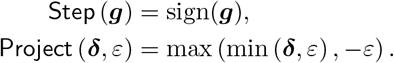

For *l*_2_-norm constraints, `Step`(·) normalizes the gradient (so that ∥ *g*∥ _2_ = 1) and `Project`(·, ·) ensures that the *l*_2_-norm of the perturbation does not exceed *ε*:

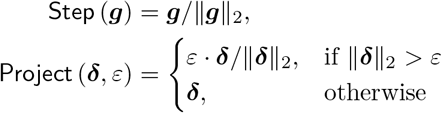

`Step`(·) and `Project`(·, ·) for *l*_1_-norm constraints are similarly defined to those of *l*_2_-norm constraints (by replacing ∥ ·∥_2_ with ∥ ·∥_1_).

## Acknowledgements

We are grateful to Cadena et al. [13] for publicly releasing their macaque V1 neural response dataset. We thank Tyler Bonnen for helpful comments on the manuscript. N.C.L.K. is indebted to the late Matthew Brennan for discussions during the early phase of this project. N.C.L.K. is supported by the Stanford University Ric Weiland Graduate Fellowship. E.M. is supported by a National Science Foundation Graduate Research Fellowship. J.L.G. acknowledges the generous support of Research to Prevent Blindness and Lions Club International Foundation (https://www.rpbusa.org/rpb/low-vision/). A.M.N. is supported by the Stanford Institute for Human Centered Artificial Intelligence.

